# Light-Sheet Fluorescence Microscopy with Scanning Non-diffracting Beams

**DOI:** 10.1101/837328

**Authors:** Hosein Kafian, Meelad Lalenejad, Sahar Moradi-Mehr, Shiva Akbari Birgani, Daryoush Abdollahpour

**Affiliations:** Department of Physics, Institute for Advanced Studies in Basic Sciences (IASBS), Zanjan 45137-66731, Iran; Department of Biology, Institute for Advanced Studies in Basic Sciences (IASBS), Zanjan 45137-66731, Iran; Optics Research Center, Institute for Advanced Studies in Basic Sciences (IASBS), Zanjan 45137-66731, Iran

**Author notes:** Correspondence shall be addressed to Daryoush Abdol-lahpour via.

## Abstract

Light-sheet fluorescence microscopy (LSFM) has now become a unique technique in different fields ranging from three-dimensional (3D) tissue imaging to real-time functional imaging of neuronal activities. Nevertheless, obtaining high-quality artifact-free images from large, dense and inhomogeneous samples is the main challenge of the method that still needs to be adequately addressed. Here, we demonstrate significant enhancement of LSFM image qualities by using scanning non-diffracting illuminating beams, both through experimental and numerical investigations. The effect of static and scanning illumination with several beams are analyzed and compared, and it is shown that scanning 2D Airy light sheet is minimally affected by the artifacts, and provide higher contrasts and uniform resolution over a wide field-of-view, due to its reduced spatial coherence, self-healing feature and higher penetration depth. Further, the capabilities of the illumination scheme is utilized for both single and double wavelength 3D imaging of a large and dense mammospheres of cancer tumor cells as complex inhomogeneous biological samples.

---

Light-sheet fluorescence microscopy (LSFM) as a novel noninvasive method provides the possibility of in-vivo volumetric imaging from the depth of thick samples, while retaining high spatial and temporal resolution simultaneously, with a minimized photodamage and phototoxicity [1]. LSFM is rapidly becoming a unique technique for functional neuroimaging in large networks [2–4], and presents the possibility of morphological imaging of a whole cleared mammalian brain, with sub-cellular resolution [5]. In contrary to the conventional wide-field and confocal fluorescence microscopes, in LSFM instead of illuminating a tiny region around the focal plane a wide layer of the sample is illuminated, and emitted fluorescence from the whole illuminated plane is captured all at once. Moreover, utilization of two separate objectives for illumination and detection results in decoupled axial and transverse resolutions, and therefore, enables the capability of three-dimensional (3D) imaging of various large and tiny specimens such as animal embryos [6], a whole mouse brain [5], and neuronal activity in a single neuron [7]. Moreover, the method is used for developmental studies in biology [8] and also for clinical histopathology of large specimens [9]. However, it should be noted that spreading of the light sheet and its deterioration, due to scattering and absorption, severely limit the capability of imaging large and dense specimens. For instance, to achieve a micron-scale axial resolution, a conventional Gaussian illuminating beam has to be focused to a micron-thick light sheet, leading to a profound action of diffraction and faster spreading of the light sheet around the confocal region. Thus, the optimum axial resolution will be limited only to a few microns in the transverse plane around the focus within the confocal range, leading to a blurry image around the focus. Furthermore, in the process of illuminating a sample with weakly absorbing or inhomogeneous structures, the light sheet is severally distributed by the emergence of a stripe pattern caused by the interference of various parts of the excitation beam after passing through inhomogeneities or absorbing structures in the sample. This effect leads to non-uniform illumination of the sample and results in artificial structures, also known as “ghost image”, in the captured images [10]. As a matter of fact, this effect is a crucial challenge in LSFM and therefore several approaches have been suggested for diminishing the issue. Rotating the sample for capturing images with different illumination directions [6], simultaneous illumination from opposite directions [11, 12], and using virtual light-sheets, formed by scanning the illuminating laser beam in a plane perpendicular to the detection direction, were initially proposed for this purpose. The first two approaches although lead to generations of uniform light-sheet in a wider field-of-view (FOV) and mitigates the ghost image, inevitably result in longer exposure of the sample to exciting illumination and therefore may induce photodamage, and also have rather complex experimental arrangements. Using non-diffracting beams such as Bessel beam [13–15], modified Bessel beam [16], and Airy beams [17–19] have also been proposed for this purpose. Although, the self-healing property of the non-diffracting beams is expected to enhance the uniformity of the illumination light sheet, their mutli-lobe intensity structure leads to simultaneous illumination of various depths, and reduces the image contrast. Therefore, finding an optimum illumination scheme for achieving high quality images with minimized artifacts, and also an enhanced resolution and contrast over a wide FOV, in large and thick samples is a severe issue to be addressed concretely.

Here, we experimentally and numerically investigate the role of the illumination on the quality of the images captured by light-sheet microscopy. We analyze and compare the LSFM images with several static and scanning light-sheets formed by different beams such as Gaussian, Bessel and Airy beams. Our results clearly reveal that illumination by the scanning 2D Airy beam results in optimal image qualities with a minimized effect of the ghost image and high contrast. Additionally, the capability of the illumination scheme is demonstrated through volumetric imaging of large and dense tumorspheres of breast cancer cells.

## THEORETICAL BACKGROUND AND NUMERICAL INVESTIGATION

The working principal of light sheet microscopy is shown in Fig. 1. A light sheet formed by an illumination objective (IO) in the *x-z* plane, illuminates a thin layer of a specimen placed around the focal plane of the objective. The excited layer emits fluorescence that is collected by a detection objective (DO) (perpendicularly oriented with respect to the light sheet), and an image is captured from the whole illuminated layer at once. Then, the light sheet or the sample are displaced with respect to each other to capture images from different depths of the sample to create 3D images. Theoretically, if the fluorophore distribution in the sample is denoted by *f* (*x, y, z*), and the intensity distribution of a very thin illumination light sheet (at a depth of *y* = *y*_0_) is represented by *I*_ill_(*x, y* = *y*_0_*, z*), the emitted fluorescence intensity distribution can be described as *I*_f_ = *f* (*x, y, z*) *· I*_ill_, and incoherent image formation is expressed as

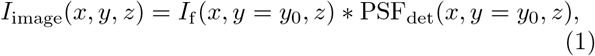

where PSF_det_ denotes the point spread function of the detection microscope, and the symbol * indicates two-dimensional convolution operation. Thus, for the very ideal case of using a thin light sheet (that is also uniform over the FOV of the detection microscope), and when the DO and IO are fixed at their position and only the sample is displaced for capturing images from various depths, the fluorophore distribution in 3D can be accurately retrieved by performing a deconvolution operation on the captured images with a given PSF_det_ at the depth of illumination. In such an ideal case the axial resolution of the LSFM is solely determined by the thickness of the light sheet, while the lateral resolution is only determined by the resolving power of the detection microscope. However, in a more realistic case, for a finite thickness of the light sheet, that is not necessarily uniform over the FOV, the axial and lateral resolutions will not be uniform over the FOV. For instance when a Gaussian beam is focused by a high numerical aperture (NA) objective, to form a thin light sheet at the waist of the illumination, the best axial resolution is only achieved within the confocal region of the focused beam and beyond that region the axial resolution is degraded due to the natural spreading of the focused beam. Similarly, the lateral resolution beyond the confocal region of the illuminating beam is also reduced since the thicker illuminated layer may lay out of the depth-of-focus of the detection microscope. Nevertheless, for a finite and uniform thickness of the light sheet across the FOV, the image formation can be described as [20]

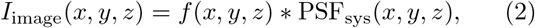

where PSF_sys_ = *I*_ill_. PSF_det_ denotes the PSF of the system that can be either numerically calculated for a given illuminating beam and a specific detection system or experimentally measured [20]. Finally, in order to retrieve the 3D fluorophore distribution, the captured image has to be deconvolved with the PSF_sys_, using an appropriate deconvultion algorithm. Therefore, a uniform resolution over the FOV can either be achieved by using a relatively thick conventional Gaussian light sheet (focused by a low-NA objective), or by using a thin light sheet formed by non-diffracting beams and their modified forms.

**FIG. 1.**
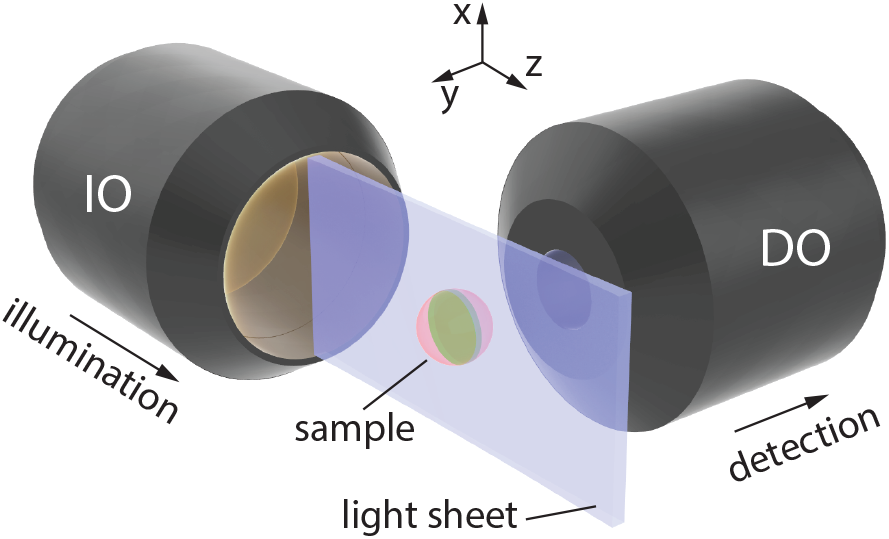
Schematics of the working principle of light sheet microscopy. Sample, located at the vicinity of the focal plane of the illumination objective (IO), is illuminated by a static or a scanning light sheet, leading to a fluorescence emission from the excited layer, whose emission is collected by the detection objective (DO), orthogonally-oriented with respect to the direction of illumination.

Additionally, the emergence of the artifacts caused by inhomogeneities in the samples is expected to be more profound when the illumination is spatially coherent (i.e. in the from of static illuminating light sheets) since the scattered beam from the inhomogeneous structures coherently superpose with the unscattered parts of the beam and lead to the generation of a modulated intensity profile in the form of stripes. On the other hand, when the light sheet is formed by scanning a beam in a transverse plane (*x-z* plane in Fig. 1) and the camera is set to integrate over several scanning cycles (to form a virtual light sheet), the area behind the obstacles are illuminated from different directions and hence the uniformity of the illumination is enhanced. Moreover, it is anticipated that the self-healing feature of the nondiffracting beams may suppress the artifacts and provide longer penetration depths of illumination in the samples that eventually lead to higher image qualities in two and three dimensions. Nevertheless, non-diffracting beams are composed of an intense main lobe accompanied by low-intensity structures that act like an energy reservoir for the main structure in a way that the power flux from the surrounding structures towards the main lobe during the propagation provides the invariant propagation and also robustness upon encountering obstacles or inhomogeneities in the propagation medium. When using these beams for creating light sheets in LSFM, the fluorescence emissions from several depth simultaneously excited by the supporting structures of the beams may also reach the detector and deteriorate the image quality. Therefore, one must consider a tradeoff between the artifacts caused by the obstacles when using conventional light sheets, on one hand, and reduction of the image contrast as a result of simultaneous illumination of various depths when using non-diffracting light sheets, on the other hand.

For a through investigation of the effect of the illuminating light sheet we conducted numerical simulations of the LSFM based on 3D beam propagation method (BPM) [21, 22] with several static and scanning illumination light sheets. Several non-fluorescent micro-beads with a diameter of 4.4 *μ*m were embedded in a uniform fluorescent medium. To simulate an inhomogeneous specimen, the refractive index of the micro-beads and the surrounding medium were selected to be different by Δ*n* = 0.2. Different illuminating beam profiles including Gaussian, Bessel, 1D and 2D Airy beams with the same widths of the main lobe (2.4 *μ*m full-width at half-maximum (FWHM), at the beam waist), similar number of side lobes for the Bessel and Airy beams, and a wavelength of *λ* = 473 nm were used for generating two static (Gaussian, and 1D Airy), and three scanning light sheets (Gaussian, Bessel and 2D Airy). The beam profiles are presented in Supplementary Figure 1. Static light sheets were created by 1D Gaussian and exponentially truncated Airy beams that are localized along *y*-axis, but uniformly extended along *x*-axis. The scanning light sheets were formed by gradually displacing the center of a Gaussian beam focused by a spherical lens and the core (main lobe) of the Bessel and 2D Airy beams along *x*-axis. Additionally, all of the beams were arranged in such a way that their waist were located in the middle of the FOV along *z*-axis (Supplementary Figure 2). The micro-beads were placed at different depths in the sample, so that the effect of simultaneous illumination of various depths by the side structures of the Bessel and Airy beams could be assessed as well. The static and scanning light sheets were propagated through the micro-beads and 3D beam propagation profiles were recorded in the same three-dimensional domain for all beams. Besides, in order to account for the fact that the spheres were not fluorescing, the 3D intensity profiles were set to zero over the volume of each sphere. Eventually, the recorded 3D intensity profiles were convolved with PSF_det_ calculated for an imaging system with NA = 0.42 at a wavelength of 515 nm based on the Born-Wolf model [23]. Calculated detection PSF is illustrated in Supplementary Figure 3. Further details of the numerical investigations are provided in the Methods section.

Simulated LSFM images of the sample for static Gaussian, static 1D Airy, and scanning Gaussian light sheets are shown in Fig. 2(a-c). Moreover, for a quantitative assessment of the severeness of the stripes after the particles, the modulus of the 2D intensity gradient, in the form of 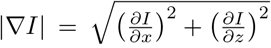, was calculated for all images after individually normalizing each image to its maximum value. Higher values of the gradient modulus would indicate stronger appearance of the stripes. The map of the intensity gradient modulus of the images are presented in Fig. 2(d-f), and their line profile, at a *z* position indicated by the dashed line, are shown in Fig. 2 (g). Obviously, the stripes are more profound in the case of the static Gaussian light sheet while their strength is relatively moderated in the case of the static 1D Airy light sheet. Nevertheless, both images are considerably affected by the presence of the stripes because of the coherence of the illumination. This can be simply verified by comparing the image associated to a scanning Gaussian light sheet (Fig. 2(c)) with the LSFM images of the static light sheets and also by comparing the intensity gradient modulus along the indicated line. It is simply seen that using a virtual light sheet is significantly enhancing the image quality in the presence of inhomogeneities.

**FIG. 2.**
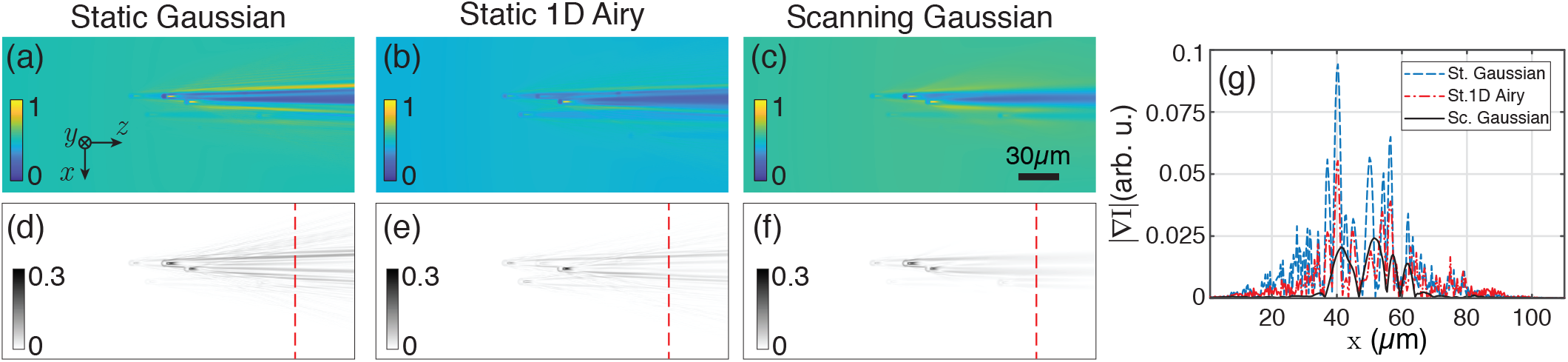
Numerically simulated LSM images of non-fluorescent micro-beads embedded in a uniform fluorescent medium, with a static Gaussian (a), static 1D Airy (b), and scanning Gaussian (c) light sheets. (d-f) corresponding intensity gradient (modulus) of (a-c); and (g) line profiles of the intensity gradient over the indicated dashed lines in (d-f) along *x*-axis. The beads are located at several depths, and the beams have identical widths at the middle of the *z*-axis.

Moreover, the simulated LSFM images of a scanning Bessel light sheet and a scanning 2D Airy light sheet are presented in Fig. 3(a, b). The modulus of the intensity gradient of the images are shown in Fig. 3(c, d), respectively. Besides, a line profile of the intensity gradient at the same *z* position are plotted in Fig. 3(e) along with that of the scanning Gaussian light sheet. Additionally, the standard deviation (SD) of the modulus of the intensity gradient for the five different light sheets over the line profiles are given in Table I. The results evidently demonstrate that the artifacts are considerably reduced by using scanning non-diffracting beams as the illumination light sheets. The highest image quality enhancement is achieved by using the scanning 2D Airy beam. Furthermore, another main factor in assessing the quality of the captured images is the contrast (visibility) of the particles with respect to the background. A qualitative inspection of the simulation results suggest that higher contrasts are achieved with static and scanning Gaussian light sheets while the lowest contrast is obtained with the scanning Bessel light sheet. This is perfectly in agreement with the previously discussed fact that the side lobes of the non-diffracting beams also contribute to image formation by simultaneously exciting different depths of the sample. Hence, the symmetric ring structure of the Bessel beam, in comparison to the Airy beam that has asymmetric distribution of the side lobes along *y*-axis, even further reduce the contrast of the images. A more rigorous quantitative comparison of the contrast of the particles will be given for the experimental LSFM images. Our simulation results vividly imply that the static light sheets although provide higher visibility for the particles, are seriously affected by the artifacts. Thus, for an experimental investigation of the quality of the LSFM images with different light sheets we limited the study to scanning light sheets.

**TABLE I.** Standard deviation of the modulus of the intensity gradient over the line profile for images with different light sheets.

| Illumination Light Sheet | SD |
| --- | --- |
| Static Gaussian | 0.016 |
| Static 1D Airy | 0.009 |
| Scanning Gaussian | 0.006 |
| Scanning Bessel | 0.003 |
| Scanning 2D Airy | 0.002 |

**FIG. 3.**
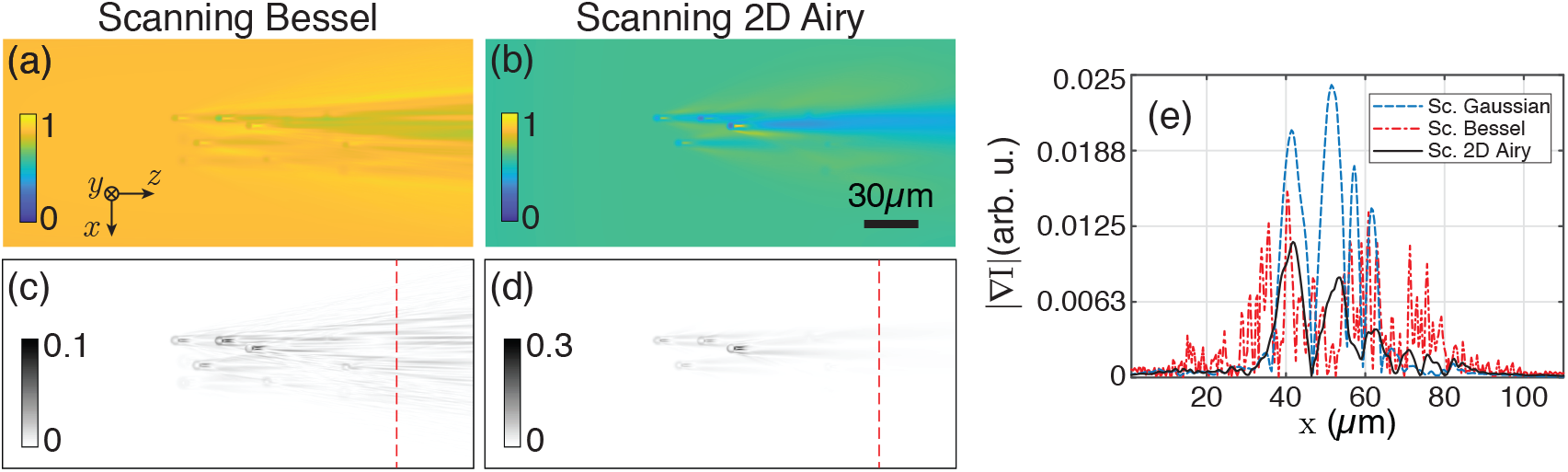
Numerically simulated LSFM images of non-fluorescent micro-beads embedded in a uniform fluorescent medium, with a scanning Bessel (a), and a scanning 2D Airy (b) light sheets.(c) and (d) present corresponding intensity gradient (modulus) of (a), and (b), respectively. (e), Line profiles of the intensity gradient over the indicated dashed lines in (c) and (d) along *x*-axis along with that of the scanning Gaussian light sheet (for comparison). The beads are located at several depths, and the Bessel and 2D Airy beams have identical widths at the middle of the *z*-axis, and a comparable number of side lobes.

## EXPERIMENTAL IMPLEMENTATION OF THE LSFM

To investigate the role of illumination in light sheet microscopy, we developed a LSFM system as illustrated in Fig. 4 and used it to image non-fluorescent micro-beads embedded in a fluorescent gel. The output of a continuous wave (CW) diode-pumped solid state laser system (*λ*_1_ = 473 nm) was collimated to an intensity FWHM of 7.5 mm, and incident on a transmissive liquid-crystal spatial light modulator (LC-SLM) with 1024 × 768 pixels, pixel pitch of 13 *μ*m and maximum phase modulation depth of *π*. An additional CW laser system with a wavelength of *λ*_2_ = 532 nm with the same FWHM could also be independently used for excitation of fluorescence when needed. The SLM was mainly used for generating various illuminating beam profiles using appropriate phase distributions. The main diffraction order of the SLM was allowed to pass an iris located at the focal plane of a lens L1, and then relayed to the scanning mirror (SM) by a 4*f* system composed of lenses L1 (*f*_1_ = 30 cm) and L2 (*f*_2_ = 15 cm). The first 4*f* system also shrinks the beam size by a factor 2 from the plane of the SLM to the plane of the SM. An additional 4*f* system composed of lenses L3 (*f*_1_ = 10 cm) and L4 (*f*_2_ = 30 cm) was used to expand and relay the beam to the back focal plane of the illumination objective (IO) (UPLAN, 10×, NA=0.3, Olympus). The lenses in both relaying systems were chosen in a way to use the full numerical aperture (NA) of the IO.

**FIG. 4.**
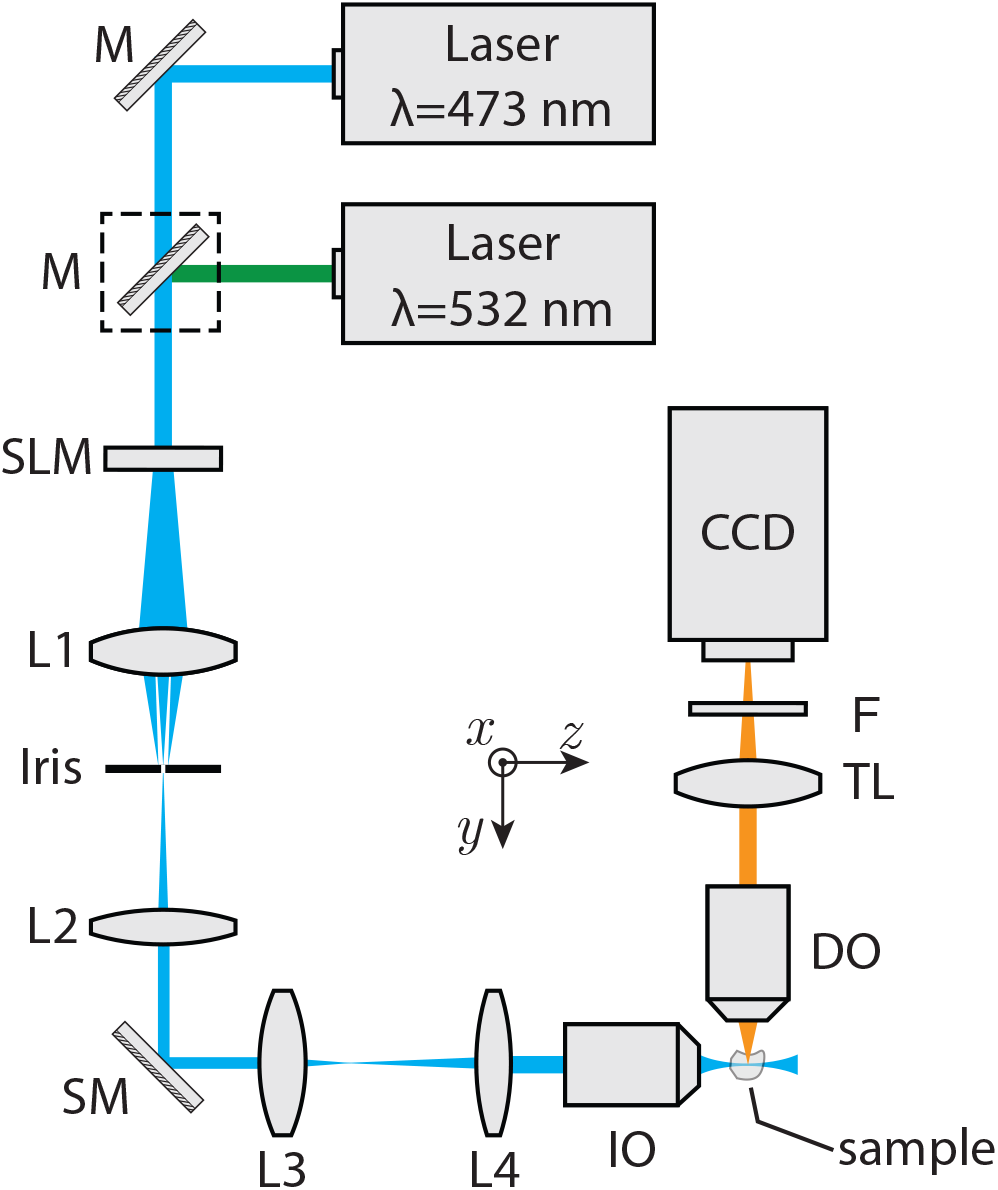
Schematic of the experimentally developed LSFM. The collimated laser beam(s) are incident on a SLM, for imprinting a desired phase profile, and then imaged into the focal plane of the illumination objective (IO) by two 4f-systems composed of the lenses L1, L2, L3, and L4. The scan mirror (SM), located at the back focal plane of the IO, provides the beam scanning capability along *x*-axis, for the generation of scanning light sheets. The waist of the light sheets overlap with the middle of the field of view (FOV) of an orthogonally oriented detection microscope, comprising a detection objective (DO), an infinity-corrected tube lens (TL), a long-pass filter (F) and a CCD camera. For excitation with 532 nm, a mirror (shown in the dashed box) was placed in front of the second laser.

Various phase profiles were prepared and imprinted on the SLM for the generation of several light sheets with different beam profiles. For the generation of a 2D Airy beam, a phase profile in the form of *α*(*x*^3^ + *y*^3^)+ *β*(*x* + *y*) was imprinted on the SLM; where, *x* and *y* are the transverse coordinates in the plane of the SLM, and *α* and *β* are arbitrary cubic and linear phase coefficients, respectively. In the case of the 2D Airy beam, the plane of the SLM was directly relayed onto the back focal plane of the IO by which a 2D spatial Fourier transformation was performed, and the beam was generated at the working distance (WD, WD = 10 mm) of the IO. On the other hand, for the generation of a Bessel beam, a conical phase profile in the from of *γρ* was imprinted on the SLM, where 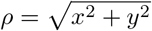 and *γ* is an arbitrary coefficient determining the width of the core of the Bessel beam. In this case, the beam was formed immediately after the SLM, and therefore an additional spherical lens, L5 (*f* = 8 cm), located 6.5 cm before the SM, was used to convert the beam to a ring-shaped beam, through a 2D Fourier transformation, and the resultant beam was then expanded by the second 4*f* system and relayed to the back focal plane of the IO and eventually transformed back to a Bessel beam at the front focal plane of the IO. Finally, for the generation of a Gaussian beam with a comparable width, the illumination scheme illustrated in Fig. 4 was modified by placing a lens L6 (*f* = 20 cm) at a distance of 4 cm before L2.

With such an arrangement, and by separately adjusting the parameters *α* and *γ*, a Gaussian beam, a Bessel beam, and a 2D Airy beam with almost identical main lobe intensity FWHMs of ~2.4 *μ*m were formed at the vicinity of the WD of the IO. The generated beam profiles (normalized to their respective maximum intensity) are depicted in Supplementary Figure 4, where it is seen that the main lobe widths of the beams and also the number of the side lobes of the Bessel and 2D Airy beams are comparable. It should be noted that the modification of the experimental setup, by adding different lenses for generating different illumination beam profiles, leads to a slight displacement of the beam formation distance from the IO (about 1 mm deviation at maximum), that was compensated by moving the object and detection microscope along the propagation direction of the illumination in each case.

Moreover, in order to form the scanning light sheets, the scanning mirror (SM) was set to oscillate at a proper frequency and amplitude to generate smooth light sheets with a dimension of 150 *μ*m along *x*-axis, and 290 *μ*m along *z*-axis. Additionally, for the case of the 2D Airy beam, the side lobes of the beam were extended along the *y*-axis and the generated light sheet had a parabolic trajectory with a maximum deviation of 3.25 *μ*m over 290 *μ*m in the sample. Moreover, the waist of the light sheets (along the propagation direction of the illumination), were adjusted to locate at the middle of the FOV of the detection microscope.

The sample was prepared by mixing and steering an aqueous suspension of nonfluorescent silica micro-beads with a diameter of 4.4 *μ*m, in a homogeneous solution of Fluorescein and gelatin. The solution was poured in a cubic transparent quartz cuvette with a dimension of (1 cm × 1 cm × 4 cm), and allowed to gradually solidify in such a way that micro-beads were firmly embedded in the fluorescent gel. The sample was mounted on a three-dimensional motorized translation stage (MT3-Z8, Thorlabs) and the emitted fluorescence from the illuminated layers of the specimen was collected and imaged by a detection microscope orthogonally oriented with respect to the illumination arm. The detection microscope was constituted of a detection objective (DO) (M PLAN 20×, APO, LWD, NA=0.42 Mitutoyo), an infinity-corrected tube lens (TTL200, Thorlabs), a longpass filter (FEL0500, Thorlabs), and a 12-bit CCD camera (GS3-U3-28S4M, PointGrey). Additionally, for the scanning light sheets, the exposure time of the CCD was adjusted in a way that multiple, but the same, number of scanning cycles of the scanning light sheets could be recorded by the camera. The translation stage and the camera were synchronized and automated to capture images from arbitrary depths inside the sample.

Normalized captured images from a depth of 100 *μ*m inside the specimen, underneath the surface facing the DO, with different illuminating scanning light sheets are depicted in Fig. 5 (a-c). Even without a quantitative assessment of the images it is obvious that the stripes are suppressed in the images with scanning non-diffracting light sheets Fig. 5(b,c), while the image with the scanning Gaussian beam is seriously deteriorated by the presence of the stripes, although in this case the particles appear with a higher contrast in comparison to the images with scanning non-diffracting beams. For a quantitative assessment of the contrast in the images we calculated the local contrast for some of the beads that were not shadowed by the presence of the stripes caused by the particles located at shorter propagation distances in the illumination direction. The selected particles are labelled with numbers 1-3, in Fig. 5(a). An 81 *μ*m^2^ square-shape region around each particle, without inclusion of the corresponding stripes (as depicted by the dashed square in Fig. 5(a)), was selected to calculate the local contrast as (*I*_max_ − *I*_min_)/(*I*_max_ + *I*_min_); where *I*_max_ and *I*_min_ are the maximum and minimum intensities in the selected region, respectively. The values of the local contrast for different illumination light sheets and different particle numbers are given in Table II. Clearly, the scanning Gaussian light sheet provides the highest contrast for most of the particles, while the lowest contrast for all three particles is achieved with the Bessel light sheet, in a prefect agreement with the simulations. Besides, the contrast values for the image with the scanning 2D Airy light sheet is comparable with that of a Gaussian light sheet, while the effect of stripes is significantly reduced in the image with 2D Airy light sheet. The modulus of the intensity gradients of the images in Fig. 5(a-c) are shown in Fig. 5(d-f). Higher values of the intensity gradient in the images imply stronger deterioration of the images by the stripes. Line profiles of the intensity gradient along *x* direction at a single *z* position, indicated by the dashed lines in Fig. 5(d-f), are plotted in Fig. 5 (g) for all light sheets, and the SD of the intensity gradient over the indicated line are given in Table III. It is seen that the stripe artifacts are significantly reduced in the image with scanning 2D Airy light sheet.

**TABLE II.** Local contrast of the micro-beads with different illumination light sheets.

| Illumination Light Sheet | Particle 1 | Particle 2 | Particle 3 |
| --- | --- | --- | --- |
| Scanning Gaussian | 0.43 | 0.41 | 0.59 |
| Scanning Bessel | 0.08 | 0.10 | 0.14 |
| Scanning 2D Airy | 0.35 | 0.43 | 0.45 |

**TABLE III.**
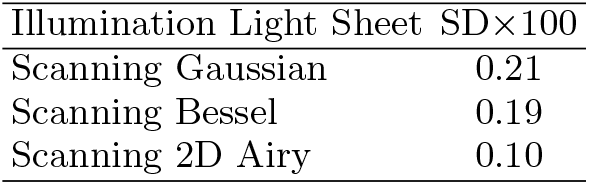
Standard deviation of the modulus of the intensity gradient over the line profile.

**FIG. 5.**
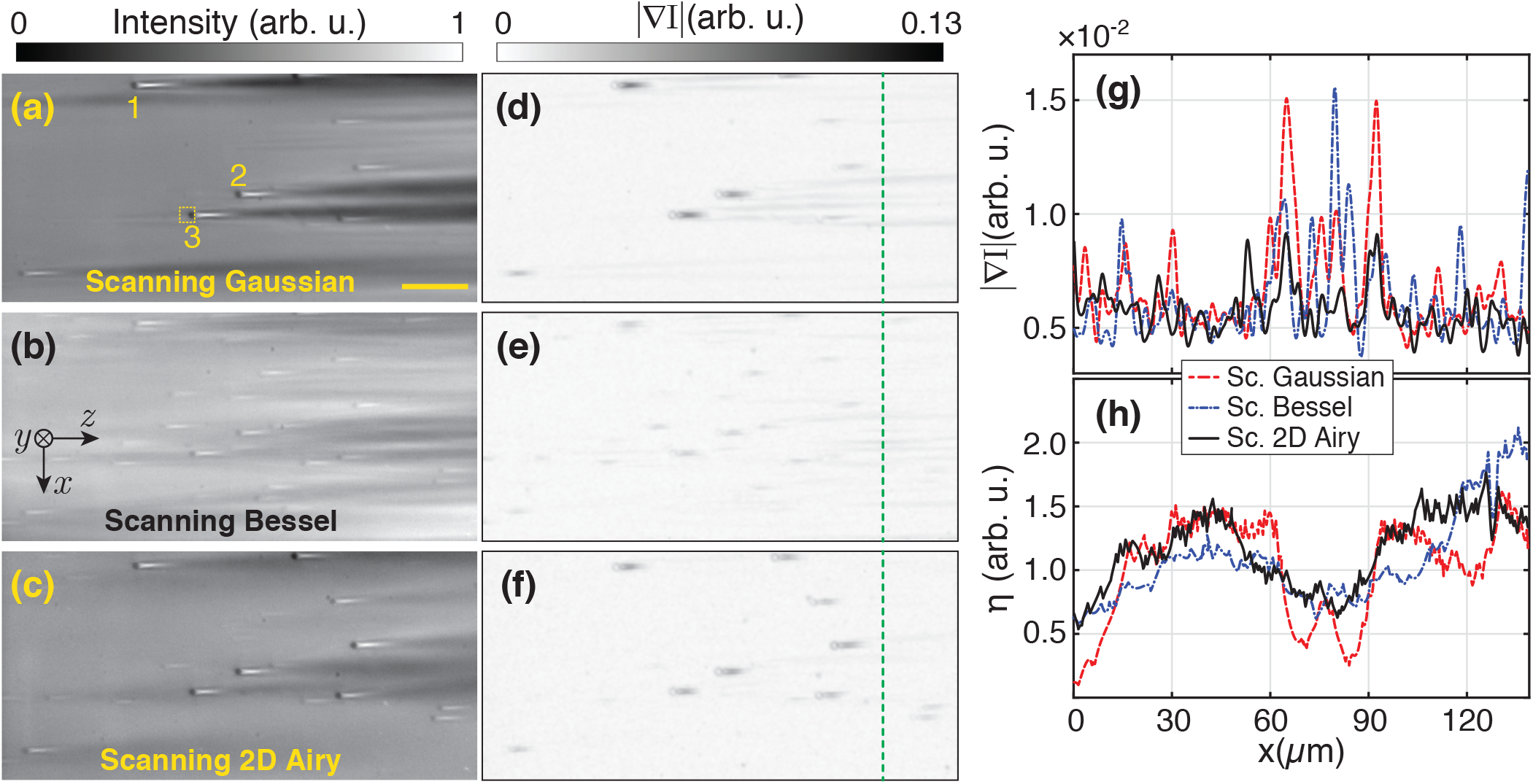
Experimentally captured LSFM images of nonfluorescent 4.4 *μ*m silica micro-beads embedded in a fluorescent gel with the scanning Gaussian (a), scanning Bessel (b), and scanning 2D Airy (c) light sheets. (d-f): Modulus of the intensity gradient associated to (a-c), respectively. (h) A line profile along *x* at a *z* position indicated by the dashed line in (d-f). (g) plot of the final to initial intensity ratio along *x*-axis.

Furthermore, the ratio of the fluorescence intensity at the final propagation distance to that of the initial propagation distance, as 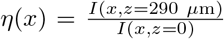, is selected as measure of the penetration depth of the light sheets in the specimen. It should be noted that the waist of all three light sheets were adjusted to be located at the middle of the FOV of the detection microscope at *z* = 145 *μ*m. In an ideal case, when the light sheet is uniform within the sample, the ratio should have minimized fluctuation around unit. For a comparison, the ratio *η*(*x*) is plotted for images with different light sheets in Fig. 5(h). Additionally, SDs of the final to initial intensity ratio in the images with different illumination light sheets are given in Table IV. These data imply that the light sheets with scanning non-diffracting beams, specially the scanning 2D Airy light sheet is more robust during the propagation in an inhomogeneous medium. The capability of resisting against the effects of the inhomogeneities and absorbing structures in dense specimen, along with providing higher contrast in the images of the structures, indicate that the scanning 2D Airy light sheet is the best illumination option in LSFM.

**TABLE IV.** Standard deviation of final to initial intensity ratio in the images with different illumination light sheets.

| Illumination Light Sheet | SD |
| --- | --- |
| Scanning Gaussian | 0.39 |
| Scanning Bessel | 0.35 |
| Scanning 2D Airy | 0.28 |

Further, we used the scanning 2D Airy light sheet in 3D LSFM imaging of mammospheres of human breast cancer cell line, as real, dense and inhomogeneous biological samples. For the sample preparation, the human breast cancer cell line MCF7 was cultured at 37°C in RPMI 1640 (Sigma-Aldrich) containing 10% Fetal bovine serum (Gibco), and 1% penicillin/streptomycin (Gibco). After the cells were 80% confluent, they were utilized for subsequent mammosphere formation. To create the mammospheres, the cells were detached and singled by adding 0.025% trypsin for five minute and then neutralized using complete media. 2 × 10^3^ cells were seeded in each well of a 96-well plate coated with agarose. 100 *μ*L of complete media was added to each well and it was changed every 2 days. Mammospheres were generated and allowed to form up to an average diameter of 200 *μ*m at day 5, and then they were harvested. The selected mammospheres were stained using Acridine orange (AO), a nucleic acid-selective fluorescent dye with an excitation maximum at 502 nm and an emission maximum at 525 nm (green) to monitor the nuclei of mammosphere-forming cells. Although the fluorescence excitation was not performed at 502 nm, but an efficient excitation was achieved with an excitation at 473 nm. The mammosphere were then immersed in a quartz cuvette (with dimensions of 1 × 1 × 4 cm^3^, and wall thickness of 1 mm) filled with a 1 mL of agarose gel, in such a manner that the mammosphere was held 150 *μ*m away from a corner of the cuvette facing the IO and the DO. The cuvette was placed in a water chamber with quartz windows, after the solidification of the agarose gel. A side of the chamber was placed adjacent to the IO (without a gap), and the other side was placed at a distance of 17.6 mm from the DO.

A 3D image stack of the mammosphere was captured with an exciting laser at a wavelength of 473 nm in the form of scanning 2D Airy light sheet with a thickness (FWHM) of 2.8 *μ*m. The detection and also the illuminating light sheet were fixed at their position, while the sample-containing cuvette was mounted on the motorized stage and displaced along −*y*-direction, to illuminate and image various depths of the sample. The motorized stage and the camera were synched in a way that after each step of displacement (by 200 nm), an image could be automatically captured. A long-pass filter with a cutoff wavelength of 500 nm was used to block the scattered laser beam. Finally, a home-developed deconvolution program based on Richardson-Lucy algorithm [24–26] was used for deconvolution of the experimentally captured images with the illumination and detection PSFs. The illumination PSF was created by directly imaging the illumination beam profile along its propagation direction at several propagation distances with a step size of 800 nm, while the detection PSF was numerically calculated according to the Gibson-Lanny model [27]. The point spread function of the whole system (PSF_sys_) was generated by multiplication of the detection and illumination PSFs and neglecting the minor deviation of 2D Airy beam from a straight line, as a result of its parabolic trajectory. The deconvolution process was performed on the 3D stack of images, first on *x-z* planes, and next on *y-z* planes. The principle of the deconvolution algorithm along with relations for the detection PSF are given in the Methods section. Deconvolved cross-sectional (*x-z*) images from three depths in the mammosphere are shown in Fig. 6(a-c). These images are presented with a green colormap to resemble the color of fluorescence emission with a peak at 525 nm, although the captured images are monochromatic. Furthermore, a video of the maximum projection of the deconvolved 3D image stack of the sample, in which the mammosphere is rotated about *x*-axis is presented as a Supplementary Video. It is easily seen that the 3D image is indeed of very high quality, without artifacts, and not affected by the side lobes of the illuminating 2D Airy light sheet.

**FIG. 6.**
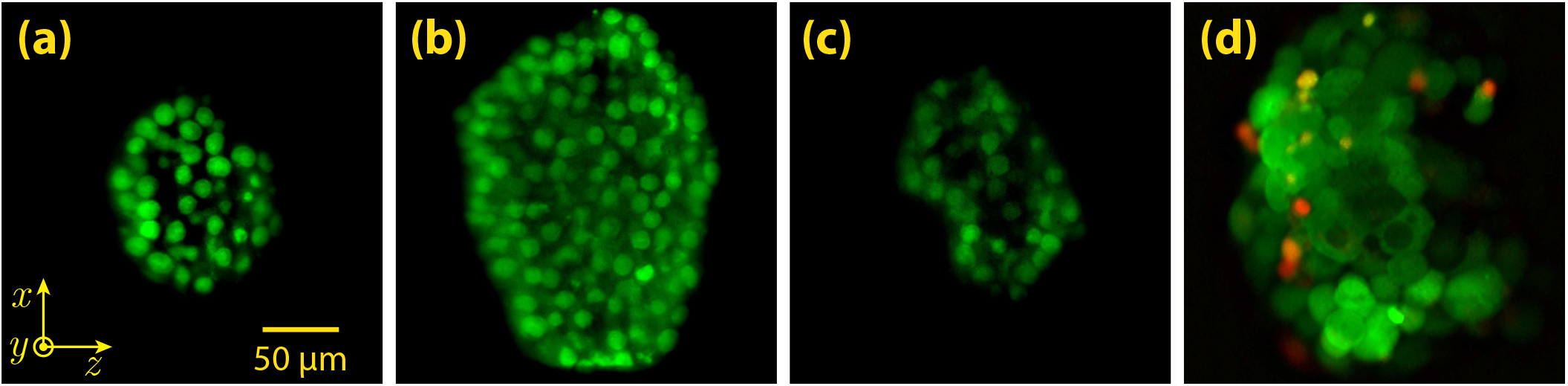
Experimentally captured cross-sectional LSFM images of a mammosphere, from three depths of 20, 120 and 220 *μ*m (a-c), respectively. The mammosphere was illuminated by a scanning 2D Airy light sheet at a wavelength of 473 nm. (d): Experimentally captured LSFM image from a depth of 100 *μm* inside another mammosphere stained with two different dyes, which differently stain live and dead cells. In this case, the mammosphere was illuminated by scanning 2D Airy light sheets of similar widths but two different wavelengths of 473 and 532 nm, separately. For each illumination wavelength an image was captured, and the images were eventually merged. Here, the red color represents the emission solely from dead cells (excitation wavelength of 532 nm), while the green color originates from live cells (excitation wavelength of 473 nm).

Furthermore, another mammosphere was prepared as described earlier and stained by calcein–acetoxymethyl (Calcein-AM; Sigma–Aldrich) and propidium iodide (PI; BioLegend). Calcein-AM is a cell permeable and nonfluorescent dye which is converted to green-fluorescent calcein, after hydrolysis by cellular esterases of live cells; hence, it is applicable to detect live cells with an excitation peak at 488 nm and maximum emission at 520 nm. On the other hand, PI is a fluorescent dye with an excitation peak at 535 nm and maximal emission at 617 nm (red) which is commonly used to detect dead cells.

Scanning 2D Airy light sheets with similar thicknesses but two different wavelengths (*λ*_1_ = 473 nm and *λ*_2_ = 532 nm) were separately used to illuminate this mammosphere, in order to discriminate dead and live cells. For this purpose, each depth of the sample was imaged separately for each excitation wavelength with a similar scanning 2D Airy light sheet. Two different longpass filters with cut-off wavelengths of 500 nm and 550 nm (FEL0550, Thorlabs) were used in the detection microscope for excitation laser beams of 473 nm, and 532 nm, respectively. A cross-sectional image of the mammosphere from a depth of 100 *μ*m, with double-wavelength excitation is shown Fig. 6(d), where green and red colors represent live and dead cells, respectively. Interestingly, some cells simultaneously produce both green and red emissions which can possibly imply that they are dying. It should be noted that both images with double excitation were taken by the monochrome CCD; however, to make a distinction between the two emissions, the images are color coded and merged. Additionally, we have to note that green color in Fig. 6(a-c) represents the nuclei stained by AO whereas green color in Fig. 6(d) pertains to the cells stained by Calcein-AM.

In conclusion, the effect of illumination in LSFM imaging with several light sheets are presented and compared. It is shown that static light sheets although provide higher contrasts in 3D imaging of inhomogeneous samples, are severely deteriorated by the appearance of artifacts. On the other hand, it is shown that scanning light sheets specially in the form of non-diffracting Bessel and Airy beams provide images with reduced artifacts. Scanning 2D Airy light sheets provide very high image contrast with significantly reduced stripe artifacts over a large FOV. This is attributed to the decreased spatial coherence, self-healing feature, asymmetric intensity distribution, and longer penetration depth of scanning 2D Airy light sheets. The illumination scheme is utilized for both single and double color excitation/detection of large, dense and inhomogeneous mammospheres of human breast cancer tumors and its shown that by a proper deconvolution of the 3D image stacks, high quality, artifact-free 3D images of such complex systems can be achieved by LSFM.

## METHODS

### Numerical Simulations

For the simulations of LSFM imaging of non-fluorescent micro-beads with a refractive index of *n*_*b*_ = 1.6 embedded in a uniform fluorescing medium with a refractive index of *n*_0_ = 1.4, BPM was implemented and used. If the electric field envelope of the illuminating beam at a given propagation distance of *z* is denoted by *U* (*x, y, z*), the field envelope upon a propagation through the medium by a step size of Δ*z* is calculated as

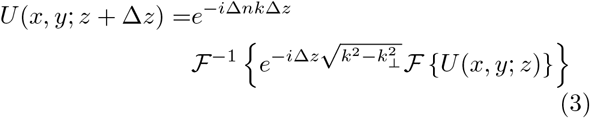

where Δ*n* = *n*_0_ − *n*_*b*_ is the difference of the refractive index between the micro-beads and the surrounding medium, *k* is the wavenumber in the surrounding medium, 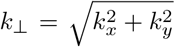 denotes the magnitude of the transverse components of the wavevector, and 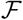, 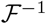 represent 2D Fourier and inverse Fourier transformations, respectively. The simulations are performed with *λ* = 473 nm, Δ*n* = 0.2 and a uniform grid spacing of Δ*x* = Δ*y* = Δ*z* = 0.2*λ*. The scanning light sheets are achieved by displacing the illuminating beams along *x*-direction by a step size of *δx* = 360 nm.

### Born-Wolf model for the detection PSF in numerical simulations of the LSFM

In the numerical investigations, the effect of the detection microscope was taken into account by performing a convolution between the simulated fluorescence emission and point spread function of the detection that was calculated based on the Born-Wolf model. According to this model, in an aberration-free system, the light intensity distribution around the focus of the detection objective, that is defined as the point-spread function of detection, PSF_det_, is given by [23]

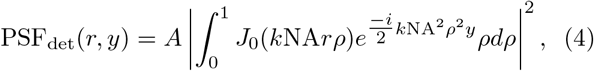

where *A* is a constant, *k* is the wavenumber of the emitted light, *J*_0_(*·*) is the zeroth-order Bessel function of the first kind, 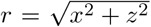 is the lateral radius on the plane of the detector (*x-z* plane), *y* is the axial distance between the focal plane and the plane of the detector, NA is the numerical aperture of the system, *ρ* = *r/a* is the radial distance normalized to the aperture radius of the objective, *a*. PSF_det_ was calculated through direct integration with NA = 0.42, *λ* = 515 nm. The transverse and longitudinal profiles of the calculated point spread function are shown in Supplementary Figure 3.

### Gibson-Lannni model for the experimental detection PSF

Unlike the Born-Wolf model, in the Gibson-Lanni model, external aberrations such as the ones originating from the index mismatch between sample, coverslip, and immersion can be taken into account. Such aberrations are characterized by optical path length difference between a ray in an ideal system and a ray under the experimental condition. According to this model, the detection point spread function is given by [27]

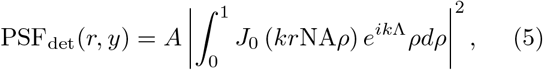

with

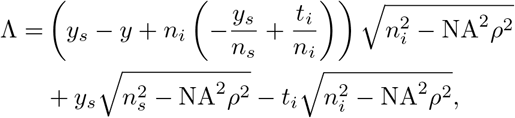

where *y*_*s*_ is the axial position of a fluorophore point source from the immersion layer, *y* is the axial distance measured from the immersion layer, *n*_*s*_ is the refractive index of the sample, *t*_*i*_ and *n*_*i*_ are the thickness and refractive index of the immersion layer, respectively. For simplicity, effects of the thin glass windows are neglected, and it is also assumed that the refractive index of the sample and that of the water are identical. The integral was directly calculated according to experimental configuration for imaging the mammospheres, with *n*_*s*_ = 1.33, *y*_*s*_ = 6.8 mm, *t*_*i*_ = 17.6 mm, *n*_*i*_ = 1, NA = 0.42, and *λ* = 525 nm. Cross-sectional distributions of the calculated PSF_det_ are presented in Supplementary Figure 5.

### Richardson-Lucy Deconvolution

With a given point spread function of the system, captured LSFM images could be properly deconvolved by using the iterative Richardson-Lucy (RL) deconvolution algorithm. Among other deconvolution approaches, the RL algorithm is shown to be capable of sufficiently retrieving the distribution even in the presence of electronic noise of the detector. The RL iterative algorithm is expressed as

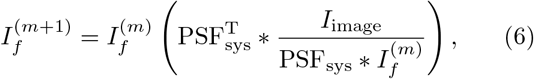

where *I*_*f*_ is the fluorophore distribution, *m* is the iteration number, *I*_image_ is the captured image, PSF_sys_ is the point spread function of the whole system, superscript T implies matrix transposition, and the symbol * indicates a 2D convolution process.

## ACKNOWLEDGMENT

The work was supported by Institute for Advanced Studies in Basic Sciences (IASBS), partially by a grant G2018IASBS12663. Additionally, D. A. specially thanks Prof. Babak Karimi, dean of IASBS, for dedicating an extra fund for the project. D. A. is also grateful to Dr. Saeed Abedinpour, Dr. Ali Ghorbanzadeh Moghaddam, Dr. Alireza Ghasemi and Dr. Hossein Fazli for partially or totally devoting their own annual research grants to this project.

## AUTHOR CONTRIBUTION

The LSFM system for this work was developed by H.K. and D.A., and H.K. performed all experiments. M.L. conducted numerical simulations and deconvolution of the experimental data. S.M-M. and S.A.B prepared the mammospheres. D.A. wrote the manuscript with contributions from all the authors, and also supervised the whole work (except for the mammosphere preparation that was supervised by S.A.B.).

## References

[1] J. Mertz, Optical sectioning microscopy with planar or structured illumination, Nat Meth 8, 811 (2011).

[2] M. B. Ahrens, M. B. Orger, D. N. Robson, J. M. Li, and P. J. Keller, Whole-brain functional imaging at cellular resolution using light-sheet microscopy, Nat Meth 10, 413 (2013).

[3] N. Vladimirov, Y. Mu, T. Kawashima, D. V. Bennett, C.-T. Yang, L. L. Looger, P. J. Keller, J. Freeman, and M. B. Ahrens, Light-sheet functional imaging in fictively behaving zebrafish, Nature Methods 11, 883 (2014).

[4] P. J. Keller, M. B. Ahrens, and J. Freeman, Light-sheet imaging for systems neuroscience, Nat Meth 12, 27 (2015).

[5] H.-U. Dodt, U. Leischner, A. Schierloh, N. Jahrling, C. P. Mauch, K. Deininger, J. M. Deussing, M. Eder, W. Zieglgansberger, and K. Becker, Ultramicroscopy: three-dimensional visualization of neuronal networks in the whole mouse brain, Nat Meth 4, 331 (2007).

[6] J. Huisken, J. Swoger, F. Del Bene, J. Wittbrodt, and E. H. K. Stelzer, Optical sectioning deep inside live embryos by selective plane illumination microscopy, Science 305, 1007 (2004).

[7] P. J. Keller and M. B. Ahrens, Visualizing whole-brain activity and development at the single-cell level using light-sheet microscopy, Neuron 85, 462 (2015).

[8] J. Huisken and D. Y. R. Stainier, Selective plane illumination microscopy techniques in developmental biology, Development 136, 1963 (2009).

[9] A. K. Glaser, N. P. Reder, Y. Chen, E. F. McCarty, C. Yin, L. Wei, Y. Wang, L. D. True, and J. T. C. Liu, Light-sheet microscopy for slide-free non-destructive pathology of large clinical specimens, Nature Biomedical Engineering 1, 0084 EP (2017).

[10] A. Rohrbach, Artifacts resulting from imaging in scattering media: a theoretical prediction, Optics Letters 34, 3041 (2009).

[11] J. Huisken and D. Y. R. Stainier, Even fluorescence excitation by multidirectional selective plane illumination microscopy (mSPIM), Optics Letters 32, 2608 (2007).

[12] U. Krzic, S. Gunther, T. E. Saunders, S. J. Streichan, and L. Hufnagel, Multiview light-sheet microscope for rapid in toto imaging, Nature Methods 9, 730 EP (2012).

[13] F. O. Fahrbach, P. Simon, and A. Rohrbach, Microscopy with self-reconstructing beams, Nat Photon 4, 780 (2010).

[14] F. O. Fahrbach and A. Rohrbach, A line scanned light-sheet microscope with phase shaped self-reconstructing beams, Optics Express 18, 24229 (2010).

[15] F. O. Fahrbach, V. Gurchenkov, K. Alessandri, P. Nassoy, and A. Rohrbach, Light-sheet microscopy in thick media using scanned bessel beams and two-photon fluorescence excitation, Optics Express 21, 13824 (2013).

[16] F. O. Fahrbach, V. Gurchenkov, K. Alessandri, P. Nassoy, and A. Rohrbach, Self-reconstructing sectioned bessel beams offer submicron optical sectioning for large fields of view in light-sheet microscopy, Optics Express 21, 11425 (2013).

[17] T. Vettenburg, H. I. C. Dalgarno, J. Nylk, C. Coll-Llado, D. E. K. Ferrier, T. Cizmar, F. J. Gunn-Moore, and K. Dholakia, Light-sheet microscopy using an Airy beam, Nat Meth 11, 541 (2014).

[18] Z. Yang, M. Prokopas, J. Nylk, C. Coll-Lladó, F. J. Gunn-Moore, D. E. K. Ferrier, T. Vettenburg, and K. Dholakia, A compact Airy beam light sheet microscope with a tilted cylindrical lens, Biomedical Optics Express 5, 3434 (2014).

[19] J. Nylk, K. McCluskey, S. Aggarwal, J. A. Tello, and K. Dholakia, Enhancement of image quality and imaging depth with Airy light-sheet microscopy in cleared and non-cleared neural tissue, Biomedical Optics Express 7, 4021 (2016).

[20] C. J. Engelbrecht and E. H. Stelzer, Resolution enhancement in a light-sheet-based microscope (SPIM), Optics Letters 31, 1477 (2006).

[21] J. A. Fleck, J. R. Morris, and M. D. Feit, Time-dependent propagation of high energy laser beams through the atmosphere, Applied physics 10, 129 (1976).

[22] M. D. Feit and J. A. Fleck, Light propagation in graded-index optical fibers, Appl. Opt. 17, 3990 (1978).

[23] M. Born and E. Wolf, Principles of Optics: Electromagnetic Theory of Propagation, Interference and Diffraction of Light (Cambridge University Press, 1999).

[24] W. H. Richardson, Bayesian-based iterative method of image restoration, Journal of the Optical Society of America 62, 55 (1972).

[25] L. B. Lucy, An iterative technique for the rectification of observed distributions, Astronomical Journal 79, 745 (1974).

[26] P. P. Mondal and A. Diaspro, Fundamentals of Fluorescence Microscopy (Springer, 2014).

[27] S. F. Gibson and F. Lanni, Experimental test of an analytical model of aberration in an oil-immersion objective lens used in three-dimensional light microscopy, Journal of the Optical Society of America A 9, 154 (1992).

