## Supplementary Figures for "Light-Sheet Fluorescence Microscopy with Scanning Non-diffracting Beams"

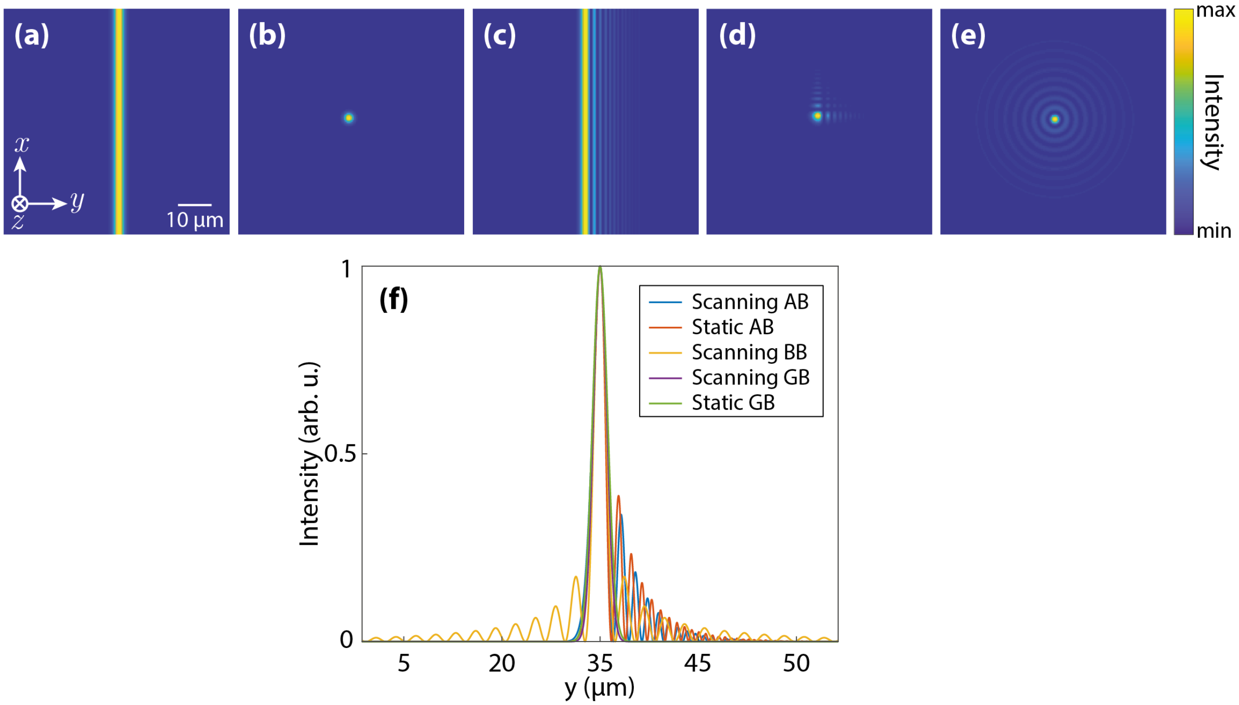


**Supplementary Figure 1 | Simulated Beam Profiles.** Transverse (x-y) normalized intensity profiles of the simulated beams at their waist for 1D Gaussian (**a**), 2D Gaussian (**b**), 1D Airy (**c**), 2D Airy (**d**), and Bessel beam. The static light sheets were formed in the x-z plane by 1D Gaussian and 1D Airy beams while the scanning light sheets were formed in the same plane by gradually displacing 2D Gaussian, 2D Airy beam and Bessel beams along x-axis. Normalized profiles of the light sheets along y-axis are shown in (**f**). Here, AB, GB, and BB indicate Airy beam, Gaussian beam and Bessel beam, respectively.

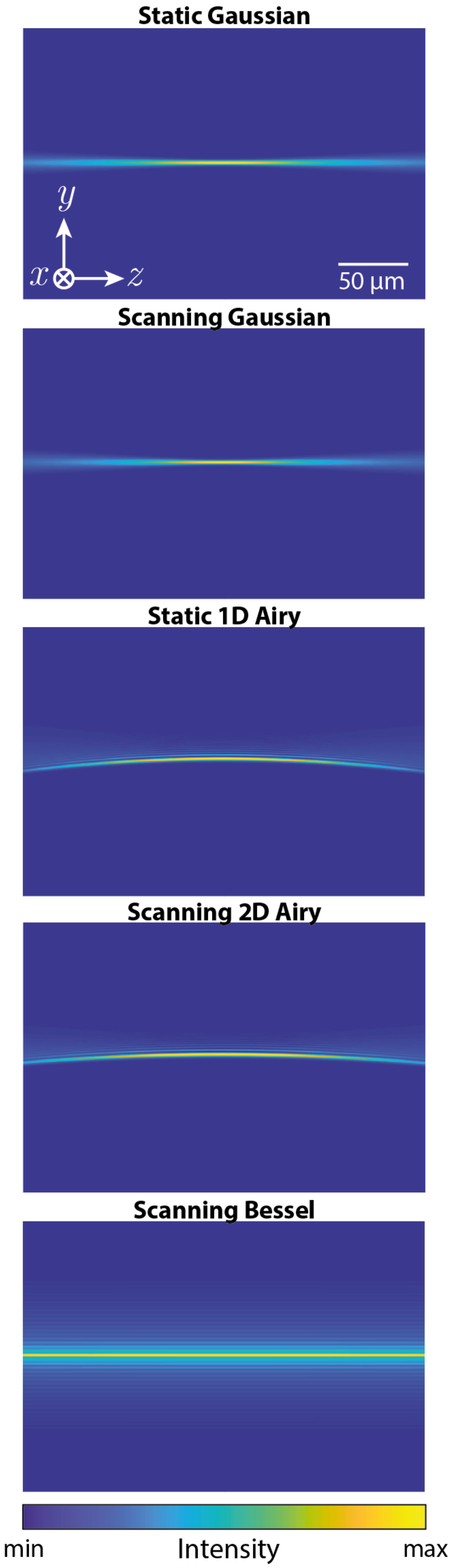


**Supplementary Figure 2 | Simulated Light-Sheet Profiles along Propagation.** Intensity profiles of the simulated light sheets along their propagation direction.

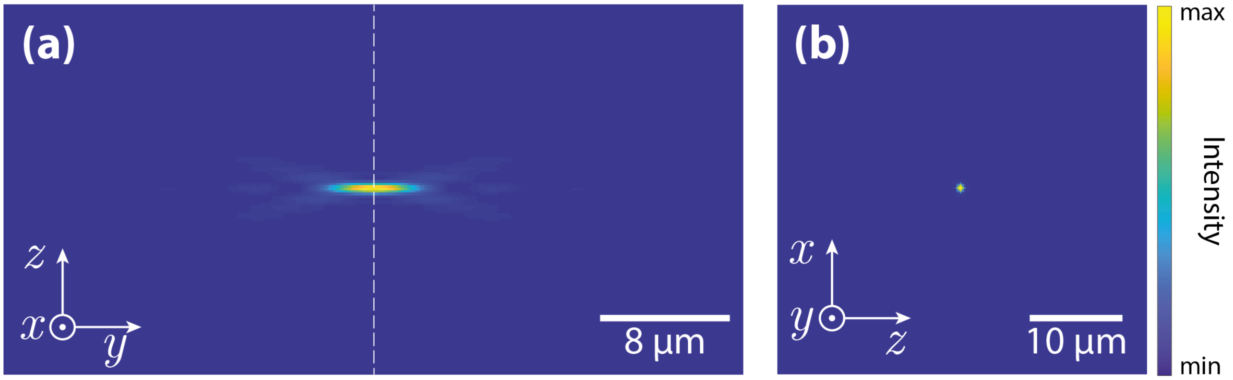


**Supplementary Figure 3 | Calculated Detection PSF based on Born-Wolf Model.** (y-z) (**a**), and (z-x) (**b**) cross-sections of the calculated detection PSF based on Born-Wolf model. The (x-z) cross-section corresponds to the focus of the PSF indicated by the dashed line in (**a**). The calculated PSF with λ=515 nm, and NA=0.42, was used to simulate the effect of the detection microscope in numerical simulations of LSFM imaging of micro-beads.

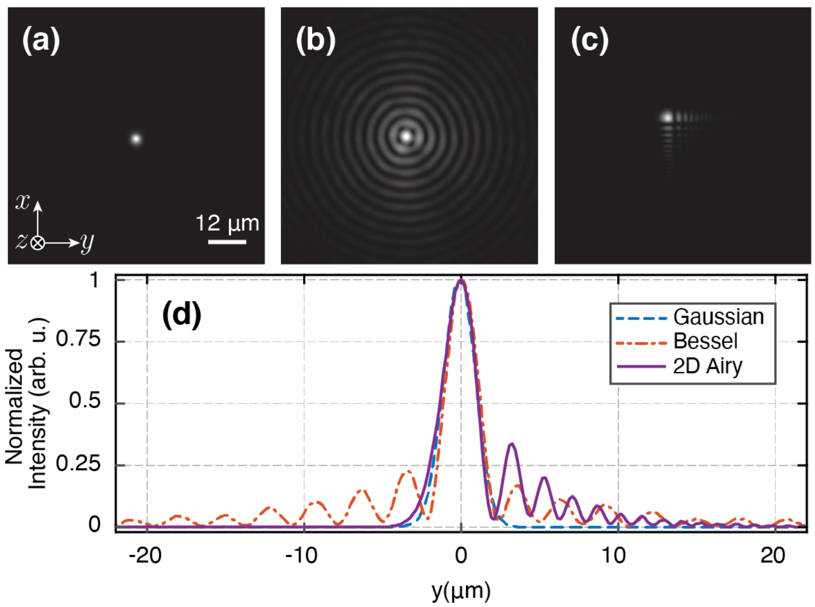


**Supplementary Figure 4 | Experimentally Generated Beam Profiles.** Transverse (x-y) normalized intensity profiles of the experimentally generated beams at their waist (i.e. at the focal plane of the IO). (**a**) 2D Gaussian, (**b**) Bessel, and (**c**) 2D Airy beams. (**e**) Normalized line profiles of the beams along y-axis.

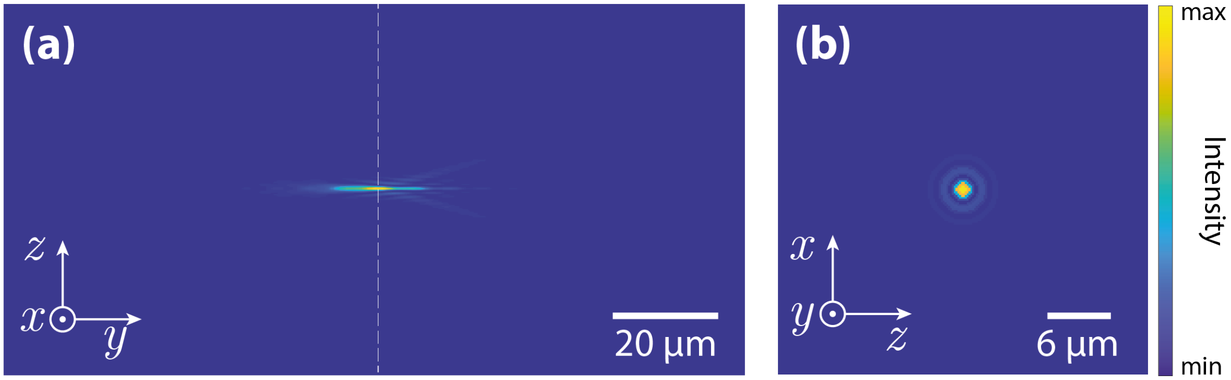


**Supplementary Figure 5 | Calculated Detection PSF based on Gibson-Lanni Model.** (y-z) (**a**), and (z-x) (**b**) cross-sections of the calculated detection PSF based on Gibson-Lanni model. The (z-x) cross-section corresponds to the focus of the PSF indicated by the dashed line in (**a**). The calculated PSF with λ=525 nm, and NA=0.42, was used to create the PSF of the whole system for deconvolution of the experimentally recorded 3D image stack of the mammospheres.
