## Supplementary figures and images for "Light-Sheet Fluorescence Microscopy with Scanning Non-diffracting Beams"

### Supplementary Video

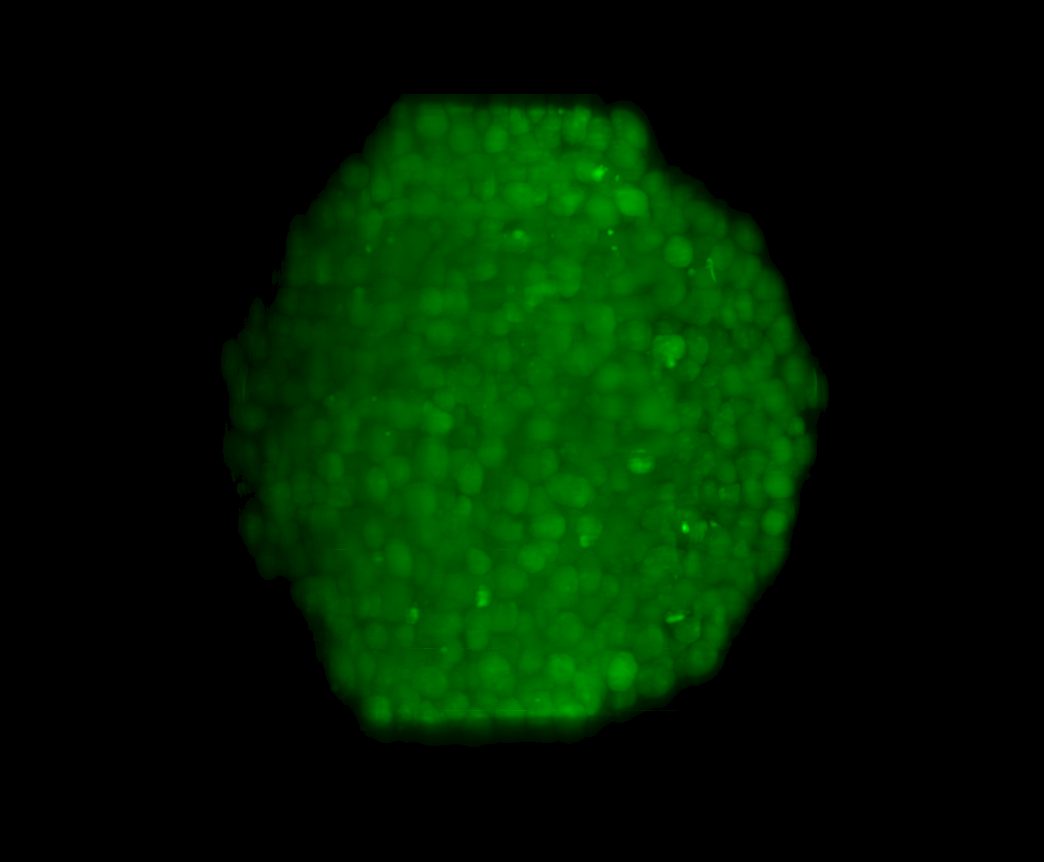
